# Superior performance of human wastewater-associated viral markers compared to bacterial markers in tropical environments

**DOI:** 10.1101/2020.10.26.355081

**Authors:** Watsawan Sangkaew, Akechai Kongprajug, Natcha Chyerochana, Warish Ahmed, Skorn Mongkolsuk, Kwanrawee Sirikanchana

**Affiliations:** Research Laboratory of Biotechnology, Chulabhorn Research Institute, 54 Kampangpetch 6 Road, Laksi, Bangkok, 10210, Thailand; CSIRO Land and Water, Ecosciences Precinct, 41 Boggo Road, Qld 4102, Australia; Center of Excellence on Environmental Health and Toxicology, CHE, Ministry of Education, 272 Rama 6 Road, Ratchathevi, Bangkok, 10400, Thailand

**Keywords:** Microbial source tracking, fecal pollution, quantitative PCR, freshwater, seawater, indicator viruses.

## Abstract

Identifying human sewage contamination via microbial source tracking (MST) marker genes has proven useful for effective water quality management worldwide; however, performance evaluations for these genes in tropical areas are limited. Therefore, this research assessed four human-associated MST marker genes in aquatic environments of Central Thailand: human polyomaviruses (JC and BK viruses [HPyVs]), bacteriophage crAssphage (CPQ_056), *Lachnospiraceae* Lachno3, and *Bacteroides* BacV6-21. HPyV and crAssphage assays were highly sensitive and specific to sewage (*n* = 19), with no cross-detection in 120 swine, cattle, chicken, duck, goat, sheep, and buffalo composite fecal samples. Lachno3 and BacV6-21 demonstrated high sensitivity but moderate specificity; however, using both markers could improve specificity to >0.80 (max value of 1.00). The most abundant markers in sewage were Lachno3 and BacV6-21 (5.42-8.02 and non-detected-8.05 log_10_ copies/100 mL), crAssphage (5.28-7.38 log_10_ copies/100 mL), and HPyVs (3.66-6.53 log_10_ copies/100 mL), respectively. HPyVs showed higher levels (up to 4.33 log_10_ copies/100 mL) and higher detection rates (92.7%) in two coastal beaches (*n* = 41) than crAssphage (up to 3.51 log_10_ copies/100 mL and 56.1%). HPyVs were also found at slightly lower levels (up to 5.10 log_10_ copies/100 mL), but at higher detection rates (92.6%), in a freshwater canal (*n* = 27) than crAssphage (up to 5.21 log_10_ copies/100 mL and 88.9%). Overall, both HPyVs and crAssphage are suggested as human sewage-associated MST markers in aquatic environments of Central Thailand. This study underlines the importance of characterizing and validating MST markers in host groups and environmental waters before including them in a water quality management toolbox.

## 1. Introduction

Fecal pollution is a significant water pollution problem, which may cause public health risks through the fecal–oral route. Human-derived pollution poses higher microbial risks than animal wastewaters due to human pathogenic waterborne microorganisms (Soller et al., 2015, 2014). Microbial source tracking (MST) is a research area studying the microorganisms excreted in feces and urine, identifying their association with their host sources, e.g., humans and animals (Ahmed et al., 2020; Kongprajug et al., 2019a; Sirikanchana et al., 2014). Pinpointing pollution sources could lead to effective pollution mitigation and improved risk assessment (Ahmed et al., 2019b; Ballesté et al., 2020; Zhang et al., 2019).

Various MST methods have been developed and validated to discriminate between contamination from different hosts. Of these, quantitative polymerase chain reaction (qPCR) is the most widely used method (Harwood et al., 2014). Among human-associated viral MST markers, human polyomaviruses (HPyVs) have been used as indicators of sewage pollution in many regions (Ahmed et al., 2019a; McQuaig et al., 2009; Rachmadi et al., 2016; Staley et al., 2012). HPyVs are nonenveloped, icosahedral, double-stranded DNA viruses, belonging to the *Polyomaviridae* family (Moens et al., 2017). HPyVs BK and JC viruses, in the *Betapolyomavirus* genus, cause nephropathy and progressive multifocal leukoencephalopathy, respectively (Moens et al., 2017). CrAssphage is a bacteriophage, which was first discovered by a comparative metagenomic approach called cross-assembly (crAss). CrAssphage has been suggested as an indicator of human fecal pollution in environmental waters (Farkas et al., 2020) and as a viral indicator of raw wastewater (Tandukar et al., 2020). Recently, crAssphage monitoring has been expanded to investigate fomite and hand contamination to potentially prevent pathogenic microbial fecal–oral transmission (Park et al., 2020).

In the USA and Australia, human-associated bacteria in the family *Lachnospiraceae* (Lachno3) and the genus *Bacteroides* (BacV6-21) have also been proposed as MST markers for human fecal pollution, and qPCR assays have recently been developed to detect them (Ahmed et al., 2019a; Feng et al., 2018; Feng and McLellan, 2019). *Lachnospiraceae* are anaerobic bacteria belonging to the order *Clostridiales*, which are prevalent in the human gut microbiota. The BacV6-21 marker was initially designed from metagenomics data to target non-fecal *Bacteroides* from the urban sewer system environment, rather than solely from human feces (Feng and McLellan, 2019). BacV6-21 showed superior sensitivity to the influents of wastewater treatment plants (WWTP-inf; *n* = 40) in the USA (Feng and McLellan, 2019). BacV6-21 was designed from sewer microorganisms and HPyVs from human urine. Therefore, both were not expected to be present in human feces. In Australia, Lachno3 and crAssphage were also found in 68.9% and 37.9% of individual human fecal samples (*n* = 29), respectively, but they were both present in 100% of WWTP-inf samples (n = 3), confirming that these markers are more suitable for tracking sewage pollution rather than individual human feces (Ahmed et al., 2019a).

Untreated wastewater is a significant source of environmental water contamination, especially in low-income countries where wastewater treatment facilities may be limited (Pollution Control Department (PCD), 2020). However, as they perform dissimilarly in diverse geographical areas (Somnark et al., 2018a), MST marker genes must be characterized before being applied in new regions. This study, therefore, investigated the performance of the two viral (HPyVs and crAssphage) and two bacterial (Lachno3 and BacV6-21) markers for human-associated MST assays in Central Thailand. The markers which performed best were further assessed for their abundance in sewage, municipal wastewater influents and effluents (WWTP-inf and WWTP-eff), and environmental water, including seawater and freshwater. Making this information available will facilitate appropriate MST marker selection for mitigating water pollution.

## 2. Materials and Methods

### 2.1 Human sewage and nonhuman fecal samples

Assay performance was assessed using archived DNA from human sewage and nonhuman fecal samples. Sewage samples (*n* = 19) were previously collected from the influent sumps of onsite wastewater treatment systems belonging to 12 hospitals, with >80 inpatient beds, in Thailand’s Bangkok, Nonthaburi, Saraburi, Samut Sakhon, Samut Prakan, and Chon Buri Provinces (Kongprajug et al., 2019b). One hundred mL of wastewater influents were stored in sterile plastic bottles containing 3.33 mL of 0.3% sodium thiosulfate. One hundred and twenty nonhuman composite fecal samples were collected by combining at least 20 individual feces samples from the same animal species, which were raised in Thailand’s Bangkok, Nakhon Pathom, Phra Nakhon Si Ayutthaya, Pathum Thani, Suphan Buri, and Chai Nat Provinces. Samples were collected from the following animal species due to their high prevalence in the area: cattle (*n* = 34), chicken (*n* = 23), and swine (*n* = 38). Samples were also taking from animals of lower prevalence—buffalo (*n* = 6), duck (*n* = 5), goat (*n* = 10), and sheep (*n* = 4)—as previously described (Somnark et al., 2018b).

### 2.2 Influent and effluent samples from wastewater treatment facilities

The abundance of MST markers was characterized in WWTP-inf and WWTP-eff samples from two municipal WWTPs in Chonburi Province. One liter of grab wastewater samples (*n* = 24) was collected over six sampling campaigns (November 2018 to May 2019) at two sampling points: one after bar screening (WWTP-inf) and the other after secondary sedimentation tank treatment (WWTP-eff). The first treatment plant (WWTP1) uses an oxidation ditch process, receiving 5,780 cu.m. of wastewater per day. The second facility (WWTP2) employs an activated sludge system, accepting 71,547 cu.m. of wastewater per day. Samples were transported on ice to the laboratory within six hours of collection. Two field blanks were collected at sampling sites as quality controls and transported to the laboratory with the other samples.

### 2.3 Seawater and freshwater samples

The abundance of MST markers was also measured in seawater and freshwater samples. Seawater samples (*n* = 42) were received from seven sampling points over two beaches in Chonburi Province (Beaches A and B) from December 2018 to August 2019. Ten liters of samples were collected at 30 cm below the water’s surface. Freshwater samples (*n* = 27) were collected from nine canal stations near an industrial estate in Rayong Province (Freshwater C) from December 2019 to March 2020. One liter of samples was taken with a water sampler at 30 cm below the water’s surface. All samples were placed in sterile plastic containers and transported to the laboratory on ice within six hours of collection. Nine field blanks were also collected at sampling sites and carried to the laboratory. Field duplicates were collected to examine the reproducibility of the laboratory protocols in the three freshwater and four seawater samples.

### 2.4 Water filtration and DNA extraction

Water samples were kept at 5°C, and fecal samples at −20°C, for no more than three days until filtration and DNA extraction were performed. One hundred mL of sewage samples were filtered with a 0.22-μm-pore-size mixed cellulose ester membrane (Merck Millipore, Billerica, MA, USA). DNA was extracted from the filtered membranes and from 150–200mg of the nonhuman fecal samples using the ZR Fecal DNA MiniPrep™ Kit (Zymo Research, Irvine, CA, USA), as previously described (Kongprajug et al., 2019b). Five-hundred mL of the WWTP-inf and WWTP-eff samples were adjusted to a pH of 3.5±0.2, followed by filtration with a 0.45-μm-pore-size HAWP membrane (Merck Millipore). Ten liters of seawater samples were preacidified as described above, and 1.5 L of each sample were separately filtered. Filtered membranes were extracted using a Quick-DNA Fecal Soil Microbe Miniprep Kit (Zymo Research, USA), and DNA extracts of the same sample were combined. One liter of freshwater samples was preacidified, and 0.5 L was separately filtered. DNA was extracted with a ZymoBIOMICS DNA Microprep Kit (Zymo Research, USA), and extracts from the same samples were mixed. The DNA concentration was measured with a NanoDrop 2000 Spectrophotometer (Thermo Scientific, USA), and the DNA was stored at −80°C until use. Thirty method blanks were analyzed by processing sterile laboratory water through all steps as quality controls.

### 2.5 qPCR assays and inhibition test

The following human-associated qPCR primers were used: HPyVs (McQuaig et al., 2009), crAssphage CPQ_056 (Stachler et al., 2017), Lachno3 (Feng et al., 2018), and BacV6-21 (Feng and McLellan, 2019) (Tables S1 and S2). The qPCR protocol was conducted according to the Minimum Information for Publication of Quantitative Real-Time PCR Experiments (MIQE) guidelines (Bustin et al., 2009). The qPCR reactions were performed with a QuantStudio™ 3 Real-Time PCR System (Applied Biosystems, Thermo Fisher Scientific, USA). For each instrumental run containing unknown samples, the DNA standard, at a concentration of 5×10^4^–5×10^5^ copies per reaction, was run in triplicate as a calibration control according to a mixed model (Kongprajug et al., 2020; Sivaganesan et al., 2010). No-template controls (NTCs) in triplicate were also included in every run. The qPCR results were analyzed using QuantStudio™ Design & Analysis Software with an automatic baseline and manual adjustment of the threshold values for the HPyV (0.025), crAssphage (0.036), Lachno3 (0.036), and BacV6-21 (0.080).

Different synthetic plasmid standards (Invitrogen, Thermo Fisher Scientific, USA), containing each marker gene, were used to develop standard curves with a mixed model method (Kongprajug et al., 2020; Sivaganesan et al., 2010). The standard curves were obtained from four replicates of individual instrumental runs, each with a triplicate of six ten-fold concentrations, ranging from 5×10^1^ to 5×10^6^ gene copies/reaction. The assay limit of detection (ALOD) was defined as the lowest concentration in copies/reactions of the ten standard replicates that showed a standard deviation of C_q_ <0.5. The assay limit of quantification (ALOQ) was considered to be the lowest concentration in copies/reactions of the target gene that could be correctly quantified—in this case, the lowest concentration in the standard curve (Haugland et al., 2010; Kongprajug et al., 2020). The method limit of quantification (MLOQ) for each sample was calculated as copies/100 mL by incorporating the sample’s filtration volume, DNA extracted volume, and DNA concentration.

Inhibition analysis was performed with the dilution method. Sewage DNA extracts were diluted to 5, 10, and 20 ng and analyzed in duplicate via the HF183/BFDrev assay as previously described (Kongprajug et al., 2020). Three dilutions (0.5, 1.0, and 2.0 μL) of templates from the WWTPs and the other environmental samples were examined with the HPyV and crAssphage assays. C_q_ values for each dilution were plotted against DNA concentration, and an R^2^ of <0.90 suggested significant inhibition. Three field blanks and three method blanks, processed with the freshwater samples, were also analyzed with HPyV and crAssphage assays, and eight field blanks and ten method blanks, examined with the seawater and municipal wastewater samples, were tested with the GenBac3 assay (Tables S1 and S2).

### 2.6 Qualitative performance measures

Human sewage from onsite treatment systems, rather than from municipal wastewater at WWTPs, was used to test the assays’ performances, thereby ensuring no stormwater contamination from nonhuman host sources. Five performance criteria were compared for each assay, with true-positive (TP) denoting the number of positive target samples, true-negative (TN) the number of negative nontarget samples, false-positive (FP) the number of positive nontarget samples, and false-negative (FN) the number of negative target samples. Sensitivity was determined by TP/(TP + FN), specificity by TN/(TN + FP), accuracy by (TP + TN)/(TP + FP + TN + FN), positive predictive value by TP/(TP + FP), and negative predictive value by TN/(TN + FN). In addition, this study selected the previously published crAssphage marker results for the archived sewage and nonhuman fecal sample sets, but the qualitative performance criteria were recalculated for comparison with the HPyV, Lachno3, and BacV6-21 markers (Kongprajug et al., 2019b).

### 2.7 Statistical analyses

Statistical analyses were carried out using an R statistical program (R Core Team, 2019). A chi-squared test of two proportions was used to compare significant differences between the assays’ performance measures. Significant differences in abundance were tested with one-way analysis of variance (ANOVA) using Holm-Sidak’s multiple comparisons. Datasets containing non-detects were analyzed via nonparametric survival analysis (Helsel, 2012). The significance of differences for datasets containing non-detects was examined via the generalized Wilcoxon test (Peto−Prentice test) for multiple comparisons, with Holm’s bias correction. A paired sample comparison for data containing non-detects was also conducted with the paired Prentice−Wilcoxon test. Correlation analysis was performed using Kendall’s Tau on U-Score rank, and water samples were grouped using principal component analysis and k-means clustering. The optimum number of clusters was determined by 30 indices using the NbClust package (Charrad et al., 2014).

## 3. Results

### 3.1 Standard curves and quality controls

The qPCR standard curves for the HPyV, crAssphage, Lachno3, and BacV6-21 markers expressed PCR efficiencies in a range of 1.943–2.019 (Table S3). The ALOD ranged from 20 copies/reaction in the crAssphage assay to 40 copies/reaction in the HPyV assay, and 50 copies/reaction in the Lachno3 and BacV6-21 assays. The ALOQ was 50 copies/reaction for all assays. In negatively detected environmental samples, the MLOQ ranged from 1.94–2.40 log_10_ copies/100 mL for HPyVs and from 1.80–3.00 log_10_ copies/100 mL for crAssphage. One negative WWTP-eff sample had an MLOQ of 3.29 log_10_ copies/100 mL.

Using the HPyV and crAssphage assays, qPCR inhibition was not observed in 24 WWTP and 41 seawater samples, nor was it found in 19 sewage samples using the HF183/BFDrev assay as previously reported (Kongprajug et al., 2020). However, four of 27 freshwater samples (14.8%) expressed inhibition when identified with the HPyV and crAssphage assays; therefore, to decrease the inhibition effect, the lowest dilution of the DNA template was used for the calculations.

Reproducibility of the laboratory protocols using the HPyV and crAssphage assays showed acceptable coefficients of variation in all representative environmental water samples (1.28–19.45%; Table S4). Seven of 63 NTCs (11.1%) were detected, but, as they were found at a late C_q_ of higher than 37.1, they could have been neglected. All field blanks and method blanks were negative, showing that no contamination occurred in field and laboratory processing steps.

### 3.2 Qualitative performance measures

All positive samples contained target genes concentrations that were higher than their corresponding ALOQ values in all assays, and the detectable-but-not-quantifiable (DNQ) data was not observed. The HPyV and crAssphage viral markers were absolute, host-specific, and sensitive; both were detected in all targeted sewage samples and absent in all nontargeted animal fecal samples (Table 1 and Table S5). The Lachno3 and BacV6-21 bacterial markers exhibited comparable sensitivity and negative predictive value to the viral markers (chi-squared test of two proportions; *p*>0.05), but they had significantly lower specificity, accuracy, and positive predictive value (*p*<0.05). One negative sewage sample in the BacV6-21 assay presented 5.90, 6.53, and 7.05 log_10_ copies/100 mL for HPyVs, crAssphage, and Lachno3, respectively.

**Table 1.** Qualitative performance measures for four human-associated MST assays

| Assay | Performance criteria |  |  |  |  |
| --- | --- | --- | --- | --- | --- |
|  | Sensitivity | Specificity | Accuracy | Positive predictive value | Negative predictive value |
| HPyV | 1.000 | 1.000 | 1.000 | 1.000 | 1.000 |
| crAssphage | 1.000 | 1.000 | 1.000 | 1.000 | 1.000 |
| Lachno3 | 1.000 | 0.593 | 0.650 | 0.284 | 1.000 |
| BacV6-21 | 0.947 | 0.721 | 0.754 | 0.367 | 0.988 |

The low specificity of the bacterial markers was found by cross-detection in all nonhuman feces—excluding duck and buffalo samples for the Lachno3 and goat and sheep samples for the BacV6-21. High cross-reaction rates were observed for the Lachno3 marker in swine (73.0%) and cattle (36.4%) and for the BacV6-21 marker in swine (52.6%), which resulted in relatively low accuracy and positive predictive values. One cattle, one chicken, and 14 swine fecal samples showed false positive results for both the Lachno3 and BacV6-21 markers. The application of both Lachno3 and BacV6-21 could improve sewage specificity to 0.855, accuracy to 0.868, and positive predictive value to 0.529.

### 3.3 Marker abundance in human sewage and nonhuman fecal samples

The abundance levels in sewage (*n* = 19) were highest for both bacterial markers, Lachno3 (5.42–8.02 log_10_ copies/100 mL) and BacV6-21 (non-detected–8.05 log_10_ copies/100 mL) (Fig. 1 and Table S6). HPyVs (3.66–6.53 log_10_ copies/100 mL) showed significantly lower abundance than crAssphage (5.28–7.38 log_10_ copies/100 mL), and both were significantly lower than the bacterial markers (paired Prentice−Wilcoxon test; *p*<0.001). The swine fecal samples showed cross-detection with the Lachno3 (*n* = 27) and BacV6-21 (*n* = 20) markers of up to 9.12 and 7.66 log_10_ copies/g feces, respectively. The chicken fecal samples presented Lachno3 (*n* = 1) and BacV6-21 (*n* = 4) markers of up to 8.21 and 8.07 log_10_ copies/g feces, respectively.

**Figure 1.**
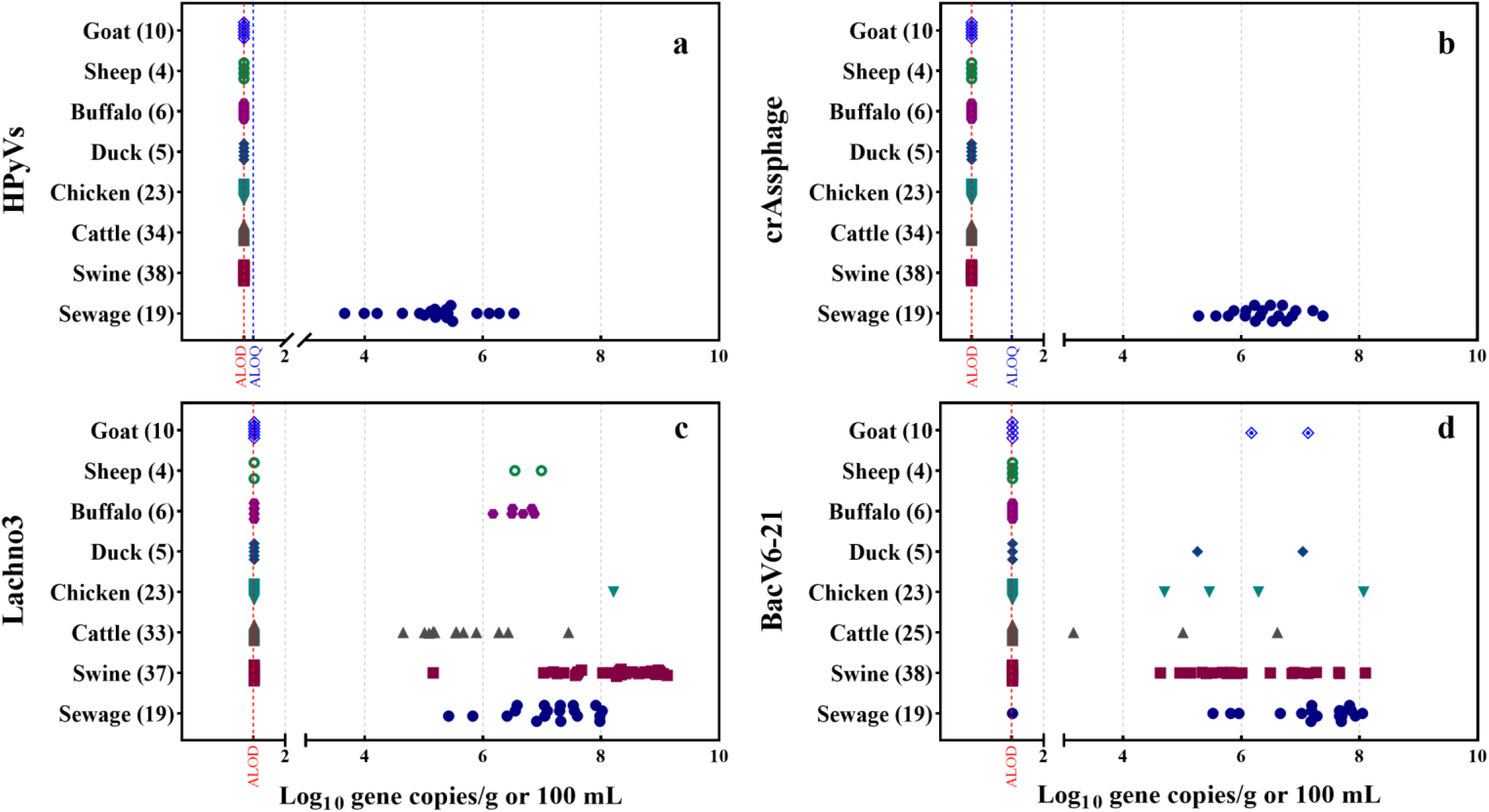
MST marker abundance in human sewage (log_10_ gene copies/100 ml) and nonhuman fecal samples (log_10_ gene copies/g): HPyVs (a), crAssphage (b), Lachno3 (c), and BacV6-21 (d). The ALOD and ALOQ are presented in log_10_ gene copies/reaction. Data for crAssphage were selected from (Kongprajug et al., 2019b) for comparison.

### 3.4 HPyVs and crAssphage in municipal wastewater

Further investigation of the highly host-associated HPyVs and crAssphage revealed a 100% positive presence in all WWTP-inf and WWTP-eff samples from the two municipal WWTPs, except for one WWTP2-eff HPyV sample (Fig. 2 and Table S7). No significant difference in HPyV abundance was observed between WWTP1-inf (4.38–6.03 log_10_ copies/100 mL) and WWTP2-inf (5.69–6.02 log_10_ copies/100 mL) and between WWTP1-eff (3.91–4.86 log_10_ copies/100 mL) and WWTP2-eff (not detectable to 5.09 log_10_ copies/100 mL) (*p*>0.05; the generalized Wilcoxon test with Holm’s bias correction). WWTP1 did not significantly remove HPyVs, with log_10_ reduction values ranging from −0.25–1.26. On the other hand, HPyV levels in WWTP2-eff were significantly lower than in WWTP2-inf (*p*<0.05; the generalized Wilcoxon test with Holm’s bias correction), with an 0.86 to a >2.40 log_10_ reduction. There were also no significant differences in HPyV levels for sewage and WWTP-inf in Thailand (*p*>0.05; one-way ANOVA with Holm-Sidak’s multiple comparisons).

**Figure 2.**
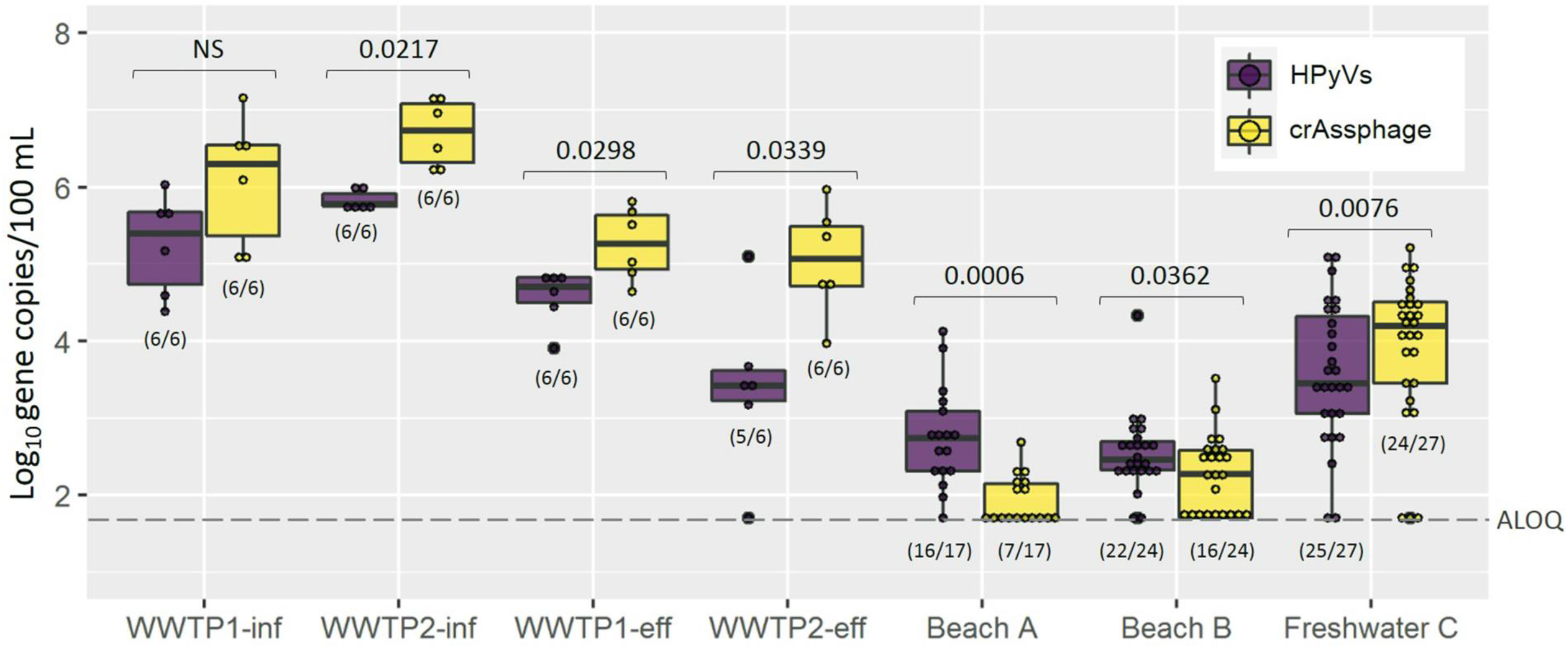
HPyV and crAssphage marker abundance in influents (WWTP1-inf and WWTP2-inf) and effluents (WWTP1-eff and WWTP2-eff) of two municipal WWTPs, seawater (Beach A and Beach B) and freshwater (Freshwater C). Box plots represent the estimated 25^th^ to 75^th^ percentiles, with the median between. Whiskers exhibit the maximum and minimum values. Numbers in parentheses describe the number of positive samples/the number of total samples. *P*-values are marked between pairs of HPyVs and crAssphage (two-tailed paired Prentice– Wilcoxon test). NS represents no significant difference. A dashed line indicates an ALOQ at 1.70 log_10_ copies/reaction.

As with HPyVs, crAssphage demonstrated similar abundance in both WWTP1-inf and WWTP2-inf (5.04–7.16 and 6.19–7.16 log_10_ copies/100 mL) and in both WWTP1-eff and WWTP2-eff (4.64–5.81 and 3.96–5.96 log_10_ copies/100 mL) (*p*>0.05; the generalized Wilcoxon test with Holm’s bias correction). WWTP1 could not reduce crAssphage (0.01– 0.96 log_10_ reduction), while WWTP2 could significantly decrease its abundance (1.15–2.52 log_10_ reduction) (*p*<0.05; the generalized Wilcoxon test with Holm’s bias correction). CrAssphage also showed no significantly different abundance levels in sewage and WWTP-inf (*p*>0.05; one-way ANOVA with Holm-Sidak’s multiple comparisons).

CrAssphage was found at significantly higher abundance levels than HPyVs in all samples except WWTP1-inf (*p*<0.05; paired Prentice–Wilcoxon test) (Fig. 2). The HPyVs and crAsssphage in municipal wastewaters were moderately correlated, showing a Kendall’s Tau value of 0.483 (*p*<0.05) (Fig. 3a).

**Figure 3.**
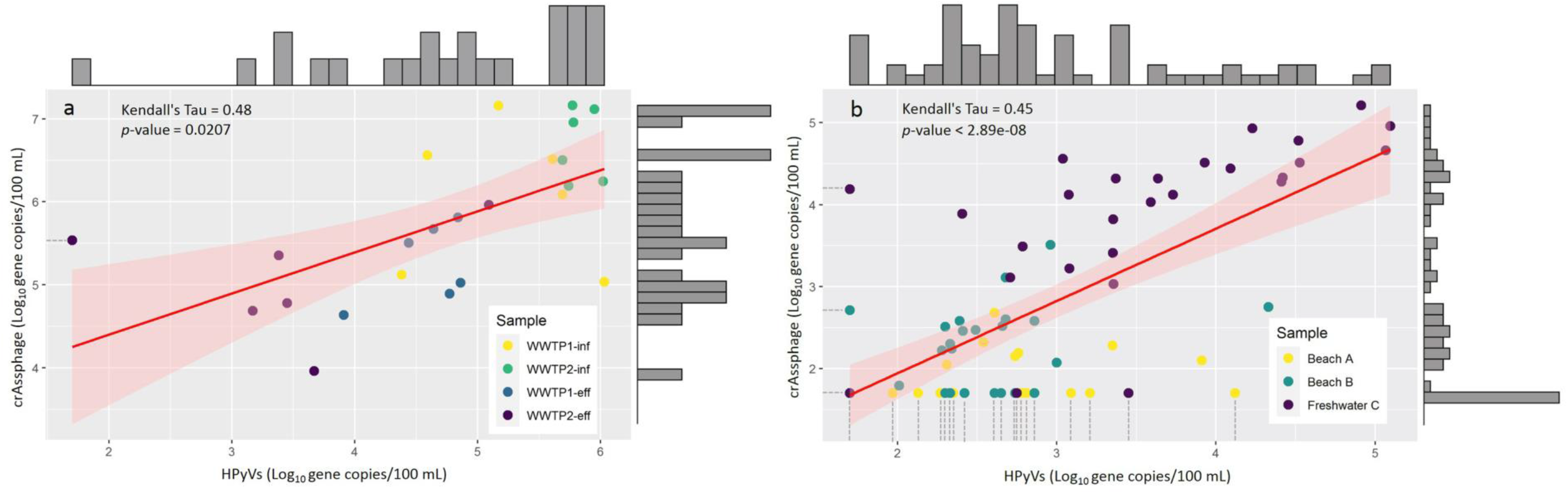
Correlation between HPyV and crAssphage markers in municipal wastewater (a) and environmental water samples (b).

### 3.5 HPyVs and crAssphage in environmental water samples

HPyVs were present in most Beach A (94.1%), Beach B (91.7%), and Freshwater C (92.6%) samples, while crAssphage was positive at a lower rate (Beach A: 41.2%; Beach B 66.7%; Freshwater C: 88.9%) (Fig. 2 and Table S7). HPyVs were found at similar abundance levels at Beaches A and B, ranging from non-detected to 4.12 log_10_ copies/100 mL and from non-detected to 4.33 log_10_ copies/100mL, respectively (*p*>0.05; the generalized Wilcoxon test with Holm’s bias correction). However, HPyV abundance was significantly higher in Freshwater C samples (non-detected to 5.10 log_10_ copies/100 mL). CrAssphage markers were significantly lower in Beach A (non-detected to 2.68 log_10_ copies/100 mL) than in Beach B samples (non-detected to 3.51 log_10_ copies/100 mL) (*p*<0.05; the generalized Wilcoxon test with Holm’s bias correction), and both beaches showed a significantly lower abundance of crAssphage than did Freshwater C samples (non-detected to 5.21 log_10_ copies/100 mL).

HPyV levels were significantly higher than crAssphage levels in Beaches A and B but lower than crAssphage in Freshwater C (*p*<0.05; paired Prentice-Wilcoxon test) (Fig. 2). In 38 HPyV-positive beach samples, 22 were also crAssphage-positive, while one HPyV-negative sample was crAssphage-positive. When using crAssphage as a screening parameter, a crAssphage-positive sample could provide 96% confidence of HPyV positivity (positive predictive value = 0.96); however, a crAssphage-negative sample could provide only 11% confidence of HPyV negativity (negative predictive value = 0.11). In freshwater samples, 23 of 25 HPyV-positive samples were also crAssphage positive, while only one HPyV-negative sample was crAssphage-positive. Therefore, crAssphage provided a positive predictive value of 0.96 and a negative predictive value of 0.33 for HPyVs. Two crAssphage-negative freshwater samples showed 2.75 and 3.45 log_10_ copies/100 mL for HPyVs, while one HPyV-negative sample contained 4.19 log_10_ copies/100 mL of crAssphage. A moderate correlation was also observed between the two markers (Kendall’s Tau = 0.453, *p*<0.001) (Fig. 3b). When separately calculated, a moderate correlation was observed for Freshwater C (Kendall’s Tau =0.558; *p*<0.05), while no significant correlation was shown for the beach samples.

Cluster analysis indicated two optimal clusters. The first dimension constituted the highest variance (80.6%) with an eigenvalue of 1.612229, while the second dimension showed a 19.4% variance with an eigenvalue of 0.387771 (Fig. 4). The two clusters placed most freshwater samples, designated as MPs, into the high contamination group, while most of the seawater samples, labeled A through G, were placed in the low contamination group. Samples were clustered into the low contamination group when (1) HPyVs were <2.78 log_10_ copies/100 mL or (2) when HPyVs were >2.78 but crAssphage was <2.60 log_10_ copies/100 mL. The distributions of both clusters were illustrated in box plots, with the high contamination group representing HPyVs from 2.79–5.10 log_10_ copies/100 mL (median: 3.82 log_10_ copies/100 mL) and crAssphage from 2.10–5.21 log_10_ copies/100 mL (median: 4.30 log_10_ copies/100 mL) (Fig. 5). The low contamination cluster contained samples with HPyV levels up to 4.12 log_10_ copies/100 mL (median: 2.54 log_10_ copies/100 mL) and crAssphage up to 4.19 log_10_ copies/100 mL (median: 2.05 log_10_ copies/100 mL).

**Figure 4.**
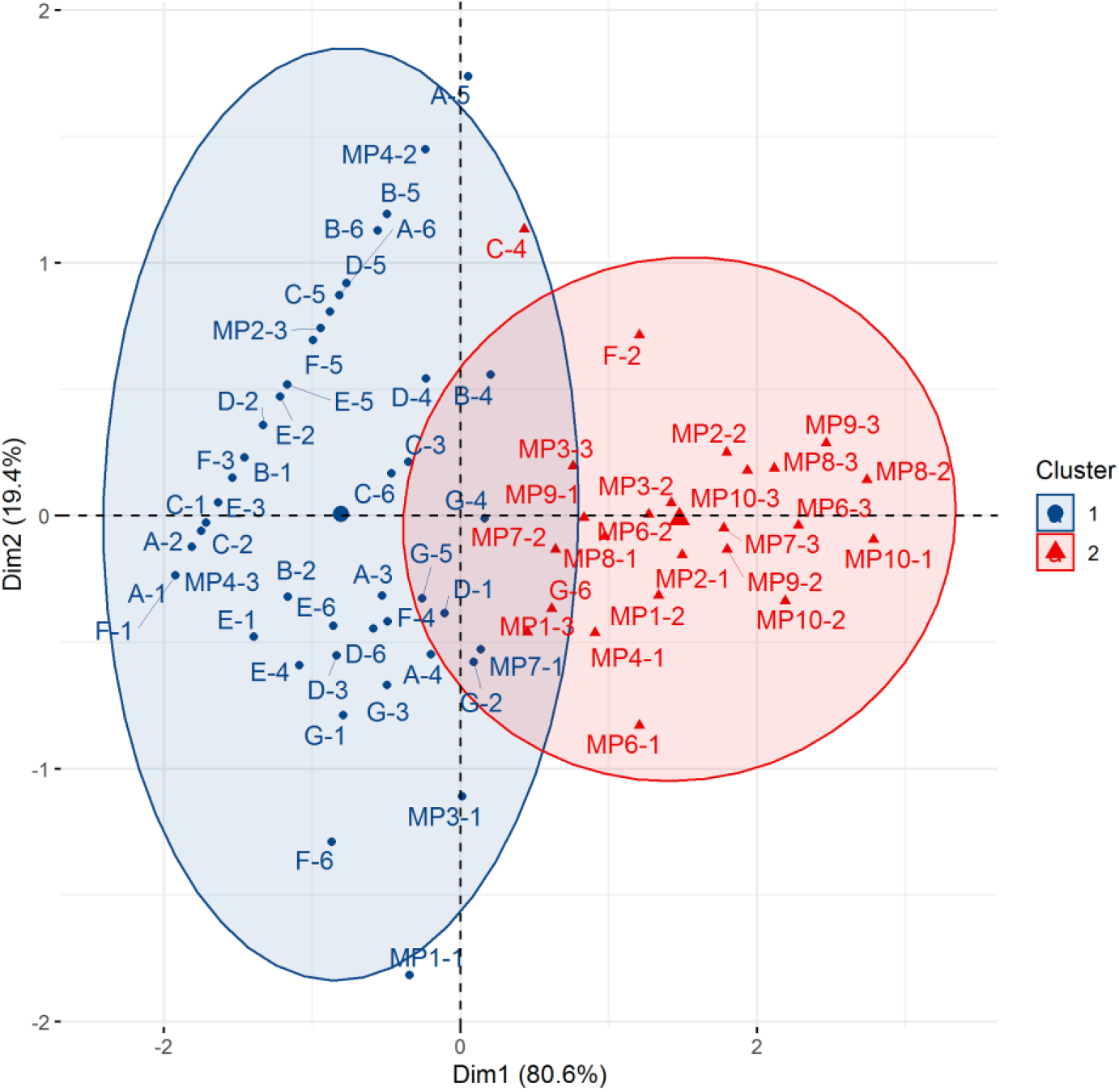
Principal component analysis (PCA) with k-mean clustering of environmental samples from Beach A (n = 17, designated as A to C), Beach B (n = 24, designated as D to G), and Freshwater C (n = 27, marked with MP).

**Figure 5.**
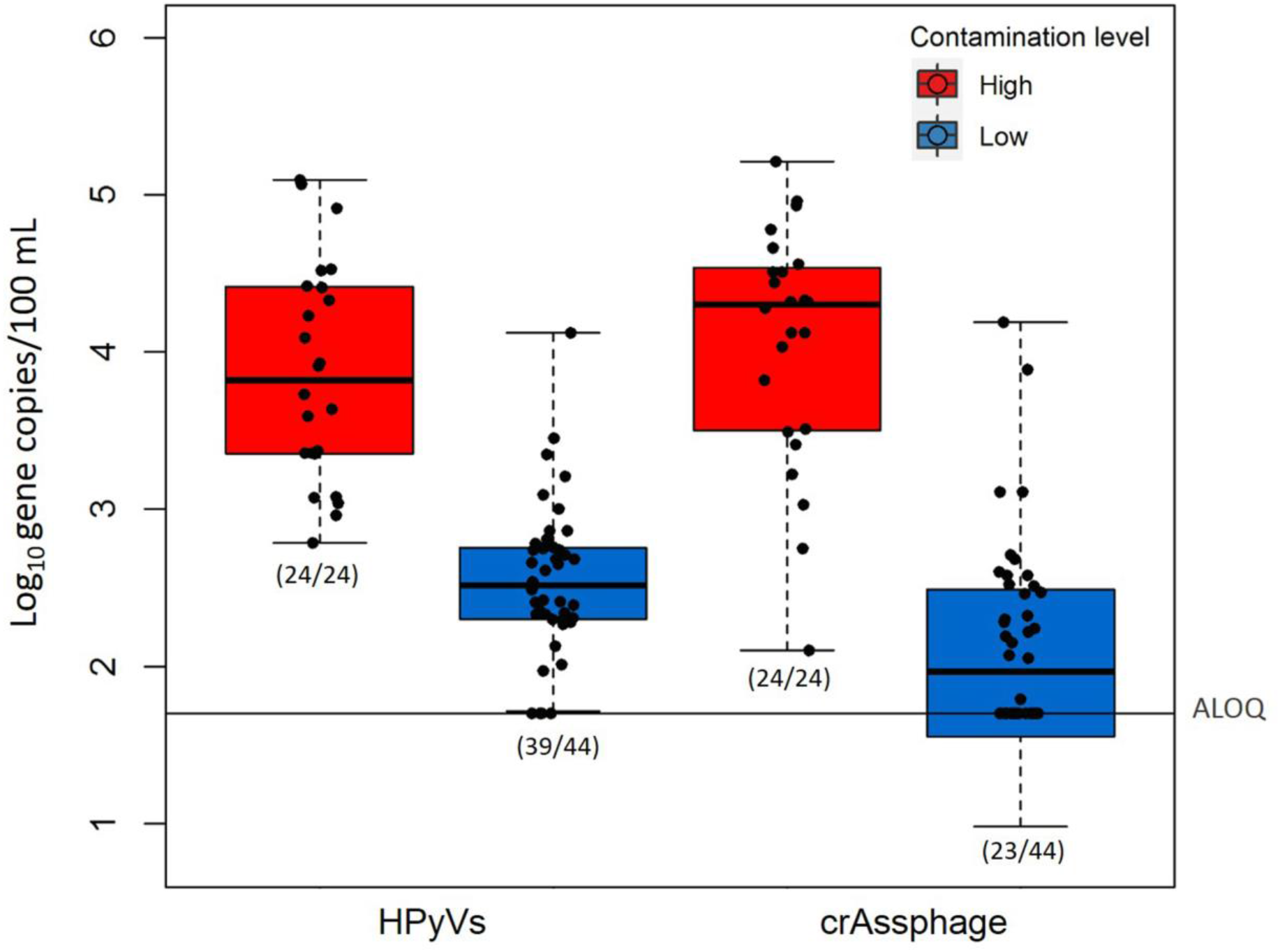
HPyV and crAssphage marker abundance in environmental samples (Beaches A and B and Freshwater C) in high and low contamination clusters. Box plots represent the estimated 25^th^ to 75^th^ percentiles, with the median between. Whiskers exhibit the maximum and minimum values. Numbers in parentheses describe the number of positive samples/the number of total samples. A dashed line indicates an ALOQ at 1.70 log_10_ copies/reaction.

## 4. Discussion

### 4.1 Higher human specificity of viral markers than of bacterial markers

The HPyV marker, targeting both BK and JC viruses, has shown 100% specificity for human sewage, with no cross-detection in any of the nonhuman fecal samples from Thailand (this study) or in other geographical regions (Ahmed et al., 2019a; McQuaig et al., 2009; Staley et al., 2012). Exceptions have been found only in one of 30 bird fecal samples and one of ten pig wastewater samples in Australia (Ahmed et al., 2019a, 2010). CrAssphage showed excellent specificity in this study; however, previous reports have indicated cross-detection in one composite swine sample from earlier work in Thailand (Kongprajug et al., 2019b) and in dog, cat, gull, poultry, and cattle samples from other geographical regions (Ahmed et al., 2018b, 2018a; Stachler et al., 2017). Overall, the results indicate that using individual viral markers— either HPyVs or crAssphage or in combination, offers the highest specificity and sensitivity and is, therefore, most appropriate for sewage detection. A similar trend showing higher specificity among viral markers than among bacterial markers has also been observed in Southeast Queensland, Australia (Ahmed et al., 2019a). This could be explained by the co-presence of certain gut bacterial genera in animals and humans (Feng et al., 2020).

Lachno3 and BacV6-21 have demonstrated relatively low specificity for human sewage detection; however, simultaneously applying both bacterial markers could improve their separate specificities to >0.80. BacV6-21 has shown high sensitivity and specificity for humans and has not cross-reacted with nonhuman fecal samples (cat, cattle, deer, dog, pig, chicken, and gull) in the USA (Feng and McLellan, 2019). However, cross-detection in cattle, chicken, pig, buffalo, and duck fecal samples in Thailand at two to eight orders of magnitude/g (this study) and in bird, cow, chicken, dog, emu, and sheep fecal samples in Australia at two to nine orders of magnitude/g have been reported (Ahmed et al., 2019a). These contradictory results emphasize geographical differences in MST marker performance. Such cross-reactions with nonhuman fecal samples have mainly obstructed bacterial marker use in agricultural regions, especially where there are similar ranges of abundance from both nonhuman samples and sewage sources. The low specificity of Lachno3 is due to its high cross-detection in cattle, pigs, sheep, and goats. Lachno3 has also been detected in nonhuman fecal samples in Australia, including bird, cat, deer, and horse at four to seven orders of magnitude/g of feces (Ahmed et al., 2019a). Originally, Lachno3 was designed from highly sewage-associated DNA regions, as identified by metagenomics data, and it demonstrated no cross-reactivity in ten cattle and nine pig samples from the Milwaukee area of the USA (Feng et al., 2018). However, cattle being raised in Thailand, with exposure to different diets and environmental stressors, may affect the animals’ gut microbiomes (Mach et al., 2020; Shanks et al., 2014, 2011). This has been supported by another marker targeting the *Lachnospiraceae* 16S V6 region, which has shown diverse prevalence in cattle with different feeding diets and across different cattle types (i.e., dairy and beef cattle) (Feng et al., 2018; Newton et al., 2011).

Other than the intrinsic presence of marker sequence in nontargeted samples, low assay specificity could also be caused by nonspecific PCR amplification (Feng et al., 2020). In a previous study, the cross-reactivity of Lachno3 with animal feces was likely caused by nonspecific qPCR amplification, as supported by an absence of Lachno3 in the metagenomic analysis (Feng et al., 2018). *Lachnospiraceae* has been reported as adversly increased nonspecific PCR amplification by lowering the annealing temperature, which could in turn increase PCR efficiency (Feng et al., 2018). In this study, the annealing temperature of the Lachno3 qPCR cycling condition was adjusted from 64°C, as originally designed, to 60°C for the following reasons: (1) the master mix reagent used in this study (i.e., 2X iTaq Universal Probes Supermix [BioRad, USA]) was recommended for use at 60°C according to the manufacturer’s instructions, and (2) higher PCR efficiency, as shown by the steeper slope of the amplification plot, was observed at 60°C compared to 64°C, according to a preliminary test (Figure S1). The corresponding PCR efficiency for the Lachno3 assay in this research was 94.3%, which is similar to that in the original study (95.5%) (Feng et al., 2018). The variation in the optimal annealing temperature between this study and the original study may have been caused by the different qPCR reagents and apparatuses (Bacich et al., 2011; Burgos et al., 2002; Nshimyimana et al., 2017; Siefring et al., 2008).

### 4.2 Higher abundance of bacterial markers than of viral markers in sewage

Though they express imperfect human source specificity, both Lachno3 and BacV6-21 demonstrated higher abundance in sewage than did the HPyVs and crAssphage viral markers. Lachno3 has been detected in WWTP-inf in the USA and Australia at five and eight orders of magnitude per 100 mL, respectively (Ahmed et al., 2019a; Feng et al., 2018). These measures are similar to the range found in Thailand. The abundance of the BacV6-21 marker in WWTP-inf has been found to be six to eight orders of magnitude per 100 mL in the USA (Feng and McLellan, 2019) and approximately eight orders of magnitude per 100 mL in Australia (Ahmed et al., 2019a)—also in a similar range to sewage in Thailand. A higher abundance of bacterial markers than of viral markers was observed in sewage and in human feces (Ahmed et al., 2019a). Although faster decay rates of bacterial than of viral DNA markers have been reported in both freshwater and seawater (Ahmed et al., 2019c; Ballesté et al., 2019; Greaves et al., 2020), the higher abundance of bacterial markers in sewage could be due to the higher excretion rates of gut bacteria than of enteric viruses or to the better adsorption of viruses to particulate matters (Horswell et al., 2010).

### 4.3 Suitability of HPyV and crAssphage as human-associated MST markers

#### 4.3.1 Higher abundance and removal of crAssphage than of HPyVs in wastewater treatment facilities

HPyVs (both BK and JC viruses) have exhibited 100% sensitivity to sewage and wastewaters, as shown in this study and in other geographical locations (Rachmadi et al., 2016); however, less sensitivity to individual urine samples has been reported (McQuaig et al., 2009). HPyVs were present in sewage and WWTP-inf in Thailand, expressed as copies/100 mL (three to six orders), at levels comparable to those detected in WWTP-inf in Australia (five to six orders) (Ahmed et al., 2019a), Japan (four orders) (Hata et al., 2013), and the USA (four to six orders) (Kitajima et al., 2014; Korajkic et al., 2020; McQuaig et al., 2009; Schmitz et al., 2016; Staley et al., 2012; Tandukar et al., 2020; Wu et al., 2020). No significant differences were observed between HPyVs from septic tanks and WWTP-inf in the USA (McQuaig et al., 2009). CrAssphage (CPQ_056) in both sewage and WWTP-inf in this study (five to seven orders, expressed as copies/100 mL) was equally abundant as that found in WWTP-inf from Indiana, USA (Wu et al., 2020), but higher than that found in WWTP-inf from elsewhere in the USA (two to four orders) (Stachler et al., 2017) and slightly lower than that found in WWTP-inf from Australia (eight orders) (Ahmed et al., 2018a, 2018b) and from two other studies in New Orleans and across the USA (up to nine orders) (Korajkic et al., 2020; Tandukar et al., 2020). CrAssphage (CPQ_056) was reported as up to six orders, expressed as copies/100 mL, in a secondary treated effluent in Louisiana, USA (Tandukar et al., 2020). Another crAssphage marker detected with a crAss-UP/LP assay, reported as four to six orders and expressed as copies/100 ml in WWTP-eff samples from Spain (Ballesté et al., 2019).

Relatively similar ranges of both HPyVs and crAssphage from different geographical regions underscore their usefulness as global sewage indicators. Neither wastewater treatment capacity nor wastewater volume received affected the concentrations of HPyVs and crAssphage, as previously stated (Crank et al., 2020). However, differences in reported quantity could result from intrinsic factors, such as different gut microbial levels due to diets and lifestyles (Honap et al., 2020; Mah et al., 2008; Turnbaugh et al., 2009), and extrinsic factors, such as temperature, sewer pipe condition, and retention time (Ahmed et al., 2019c; Okadera et al., 2020). Comparisons among different studies could also present confounding biases due to different field and laboratory protocols, analytical instruments, and data analysis methods, which could affect each virus at different levels (Agetsuma-Yanagihara et al., 2017; Crank et al., 2020; Kongprajug et al., 2020; Petcharat et al., 2020; Shanks et al., 2012).

This study presented HPyVs and crAssphage removals through wastewater treatment processes (through secondary sedimentation) of > 2.40 and 2.52 log_10_ removal, respectively, in an activated sludge treatment plant (WWTP2). The oxidation ditch treatment plant (WWTP1) removed neither viral marker. Different treatment processes could be one of the factors affecting removal efficiency, as it has been previously reported that activated sludge treatment facilities more significantly reduce crAssphage levels than do biofilter treatment plants (Farkas et al., 2019). Various maximum removal levels of HPyVs in treatment facilities at points prior to the disinfection process have been reported, ranging from 1.33 log_10_ removal in Indiana, USA (Wu et al., 2020), to 2.9 log_10_ removals in Japan and Louisiana, USA (Hata et al., 2013; Tandukar et al., 2020). CrAssphage has been removed at 2.48 log_10_ removal by a wastewater treatment facility in Indiana, USA (Wu et al., 2020).

As similarly observed in this study, crAssphage has been present in WWTP-inf at higher concentrations than HPyVs (Crank et al., 2020; Farkas et al., 2019, 2018; Tandukar et al., 2020) and has been more significantly reduced than HPyVs through wastewater treatment plants in Indiana and Louisiana, USA (Tandukar et al., 2020; Wu et al., 2020). The difference in removal efficiency between the two markers could be caused by the varying behaviors of each virus, as demonstrated by the effect concentration methods have on crAssphage but not on HPyVs (Crank et al., 2020). Moderate to high correlations between the two markers have been observed in Indiana, USA (Pearson correlation coefficient of 0.87) (Wu et al., 2020), Louisiana, USA (Pearson correlation coefficient of 0.89 and 0.91 in WWTP-inf with JC and BK, respectively) (Tandukar et al., 2020), the UK (Spearman correlation coefficient of 0.50 and 0.67 in WWTP-inf and WWTP-eff, respectively) (Farkas et al., 2019), and Italy (Spearman’s rank correlation coefficients of 0.74 and 0.85, depending on two different concentration methods) (Crank et al., 2020). Overall, the present study promotes the suitability of crAssphage as a human wastewater-associated indicator due to its consistently high abundance.

#### 4.3.2 Higher abundance in seawater and higher positive rate in freshwater of HPyVs than of crAssphage

A significantly higher abundance of crAssphage than of HPyVs in freshwater has been observed in this study, conforming to other research. In an impacted stream in Pennsylvania, USA, crAssphage (CPQ_056 assay) and HPyVs were reported at four to six and one to four orders, expressed as copies/100 mL, respectively, with no significant correlation observed (Stachler et al., 2018). Another study in the UK represented two to six orders of crAssphage CPQ_056 and up to three orders of HPyVs (JC) in freshwater, with moderate correlation (Spearman correlation coefficient of 0.49) (Farkas et al., 2019, 2018). A river in Nepal was monitored for crAssphage (CPQ_056) at four to seven orders, expressed as copies/100 mL, and HPyVs (JC and BK) of up to six orders; moderate correlation (Spearman’s rank-order correlation coefficient of 0.47) was found between crAssphage and BK, while no correlation was found between crAssphage and JC (Tandukar et al., 2018; Ward et al., 2020). However, although HPyVs were lower in abundance than crAssphage, they exhibited a higher positive rate in freshwater samples and, thus, presented higher benefits for water quality monitoring, indicating human pollution.

Although crAssphage had a significantly higher abundance in sewage than HPyVs, the present results revealed higher concentrations of HPyVs than crAssphage, with higher positive rates, in Thailand’s seawater. Similar decay rates of both markers were observed in both freshwater and seawater microcosms at 18–28°C and when exposed to sunlight (Ahmed et al., 2019c; Greaves et al., 2020). However, other physical, chemical, and biological factors could affect the fate and transport of each virus differently. In this study, no significant correlation was observed between HPyVs and crAssphage in seawater, implying the markers’ different behaviors. Further investigation is, therefore, needed to characterize factors affecting the different abundance levels in these sampling sites and to expand the investigation to other seawater and freshwater sites to develop a better understanding. In summary, this study suggests that both HPyVs and crAssphage may be used to monitor seawater and freshwater, identifying human fecal pollution in Thailand.

## 5. Conclusions

- The two viral markers, HPyVs and crAssphage, demonstrated ideal performance for human sewage source tracking in Thailand. The two bacterial markers, Lachno3 and BacV6-21, presented high sensitivity but moderate specificity, with cross-detection in poultry (chicken and duck) and livestock (swine, cattle, goat, sheep, and buffalo).
- Both bacterial markers were present in sewage at similar levels and were significantly higher than the two viral markers. CrAssphage was significantly higher than HPyVs in sewage, in the influents of WWTP2, and in the effluents of both WWTP1 and WWTP2.
- HPyVs showed higher positive rates than crAssphage in freshwater and seawater but presented significantly higher levels only in seawater. HPyVs and crAssphage expressed moderate correlations (Kendall’s Tau) in wastewater treatment and environmental waters and can be used to indicate high and low contamination clusters.
- The results suggested that HPyVs and crAssphage, or in combination, may act as human-associated MST markers in contaminated aquatic environments. This study also highlighted the necessity of validating MST markers in wastewater and local environmental waters to effectively facilitate water quality management.

## Acknowledgements

This work was financially supported by the Chulabhorn Research Institute and the Kurita Overseas Research Grant (grant no. 19Pth007 to A.K.) provided by Kurita Water and Environment Foundation.

## Appendix A. Supplementary data

Supplementary data to this article can be found online at …

