## supplementary material for "Superior performance of human wastewater-associated viral markers compared to bacterial markers in tropical environments"

**Table S1** Primers and probes used in this study

| Assay name | Primer/probe name | Primer sequence (5' - 3') | Target microorganism | Target gene | Reference |
|---|---|---|---|---|---|
| HPyV | SM2 | AGT-CTT-TAG-GGT-CTT-CTA-CCT-TT | human polyomaviruses JCV and BKV | T antigen | (McQuaig et al., 2009) |
| | P6 | GGT-GCC-AAC-CTA-TGG-AAC-AG | | | |
| | KGJ3 | FAM-TCA-TCA-CTG-GCA-AAC-AT-TAMRA | | | |
| crAssphage | CPQ_056F1 | CAG-AAG-TAC-AAA-CTC-CTA-AAA-AAC-GTA-GAG | crAssphage | Genomic region 14731−14856 | (Stachler et al., 2017) |
| | CPQ_056R1 | GAT-GAC-CAA-TAA-ACA-AGC-CAT-TAG-C | | | |
| | CPQ_056P1 | FAM-AAT-AAC-GAT-TTA-CGT-GAT-GTA-AC-TAMRA | | | |
| Lachno3 | Forward primer | CAA-CGC-GAA-GAA-CCT-TAC-CAA-A | *Lachnospiraceae* | 16s rRNA | (Feng et al., 2018) |
| | Reverse primer | CCC-AGA-GTG-CCC-ACC-TTA-AAT | | | |
| | Probe | FAM-CTC-TGA-CCG-GTC-TTT-AAT-CGG-A-BHQ1 | | | |
| BacV6-21 | Bac989f | GCT-TGA-ATT-GCA-GAG-GAA-TA | *Bacteroides* | 16s rRNA | (Feng and McLellan, 2019) |
| | Bac1162r | GCA-GTC-TCA-CTA-GAG-TCC-TCA-G | | | |
| | Bac1010p | FAM-AGT-TGA-AAG-ATT-ATG-GCC-GCA-BHQ1 | | | |
| GenBac3 | GenBac3F | GGG-GTT-CTG-AGA-GGA-AGG-T | *Bacteroidetes* | 16s rRNA | (Siefring et al., 2008) |
| | GenBac3R | CCG-TCA-TCC-TTC-ACG-CTA-CT | | | |
| | GenBacProbe | FAM-CAA-TAT-TCC-TCA-CTG-CTG-CCT-CCC-GTA-TAMRA | | | |

**Table S2** qPCR reaction mixtures and cycling conditions

| Assay name | 20-μL reaction mixtures (type A)[a] | 20-μL reaction mixtures (type B)[b] | Cycling conditions |
|---|---|---|---|
| HPyV | 0.8 μL of each 10 μM forward and reverse primers, 0.4 μL of 10 μM probe, 4 μL (40 ng equivalent) of DNA template, 4 μL of 1 μg/μL bovine serum albumin (BSA), and 10 μL of the 2X iTaq Universal Probes Supermix (BioRad, USA) | For WWTP, 0.8 μL of each 10 μM forward and reverse primers, 0.4 μL of 10 μM probe, 4 μL (40 ng equivalent) of DNA template, 4 μL of 1 μg/μL BSA, and 10 μL of the 2X iTaq Universal Probes Supermix (BioRad, USA). For environmental water, 0.8 μL of each 10 μM forward and reverse primers, 0.4 μL of 10 μM probe, 2 μL of extracted DNA, 6 μL of 1 μg/μL BSA, and 10 μL of the 2X iTaq Universal Probes Supermix (BioRad, USA) | An initial denaturation at 95°C for 3 min followed by 40 cycles of a denaturation step at 95°C for 20 s and a combined annealing and elongation step at 55°C for 1 min |
| crAssphage | 0.8 μL of each 10 μM forward and reverse primers, 0.4 μL of 10 μM probe, 5 μL (20 ng equivalent) of DNA template, 3 μL of sterile distilled water, and 10 μL of the 2X iTaq Universal Probes Supermix (BioRad, USA)[c] | For WWTP, 0.8 μL of each 10 μM forward and reverse primers, 0.4 μL of 10 μM probe, 4 μL (40 ng equivalent) of DNA template, 4 μL of 1 μg/μL BSA, and 10 μL of the 2X iTaq Universal Probes Supermix (BioRad, USA). For environmental waters, 0.8 μL of each 10 μM forward and reverse primers, 0.4 μL of 10 μM probe, 2 μL of extracted DNA, 6 μL of 1 μg/μL BSA, and 10 μL of the 2X iTaq Universal Probes Supermix (BioRad, USA). | An initial denaturation at 95°C for 3 min followed by 40 cycles of a denaturation step at 95°C for 20 s and a combined annealing and elongation step at 60°C for 1 min |
| Lachno3 | 0.8 μL of each 10 μM forward and reverse primers, 0.2 μL of 10 μM probe, 4 μL (40 ng equivalent) of DNA template, 4.2 μL of 1 μg/μL BSA, and 10 μL of the 2X iTaq Universal Probes Supermix (BioRad, USA) | NA[d] | An initial denaturation at 95°C for 3 min followed by 40 cycles of a denaturation step at 95°C for 20 s and a combined annealing and elongation step at 60°C for 1 min |
| BacV6-21 | 2.0 μL of each 10 μM forward and reverse primers, 0.2 μL of 10 μM probe, 4 μL (40 ng equivalent) of DNA template, 1.8 μL of 1 μg/μL BSA, and 10 μL of the 2X iTaq Universal Probes Supermix (BioRad, USA) | NA | An initial denaturation at 95°C for 3 min followed by 40 cycles of a denaturation step at 95°C for 20 s and a combined annealing and elongation step at 60°C for 1 min |
| GenBac3 | 0.8 μL of each 10 μM forward and reverse primers, 0.4 μL of 10 μM probe, 5 μL (20 ng equivalent) of DNA template, 3 μL of sterile distilled water, and 10 μL of the 2X iTaq Universal Probes Supermix (BioRad, USA) | 0.8 μL of each 10 μM forward and reverse primers, 0.4 μL of 10 μM probe, 2 μL of extracted DNA, 6 μL of 1 μg/μL BSA, and 10 μL of the 2X iTaq Universal Probes Supermix (BioRad, USA) | An initial denaturation at 95°C for 3 min followed by 40 cycles of a denaturation step at 95°C for 20 s and a combined annealing and elongation step at 60°C for 1 min |

[a] For human sewage and non-human fecal samples [b] For wastewater treatment plant (WWTP) and environmental samples [c] From (Kongprajug et al., 2019) [d] Not available

**Table S3** Standard curve equations and qPCR assay characteristics

| Assay name | $Slope_{std}$ | Y-intercept | PCR efficiency[a] | ROQ[b] | ALOD[c] | ALOQ[d] | $R^2$ |
|---|---|---|---|---|---|---|---|
| HPyV | -3.307 | 42.17 | 2.006 | 50 - 5 × $10^6$ | 40 | 50 | 0.998 |
| crAssphage[e] | -3.277 | 41.38 | 2.019 | 50 - 5 × $10^6$ | 20 | 50 | 0.998 |
| Lachno3 | -3.467 | 40.45 | 1.943 | 50 - 5 × $10^6$ | 50 | 50 | 1.000 |
| BacV6-21 | -3.414 | 40.70 | 1.963 | 50 - 5 × $10^6$ | 50 | 50 | 0.997 |
| GenBac3[e] | -3.296 | 39.811 | 2.011 | 50 - 5 × $10^6$ | 20 | 50 | 0.999 |

[a] PCR amplification efficiency ($E_p$) calculated by Ep = $10^{(-1/Slope)}$

[b] Range of quantification (copies/reaction)

[c] Assay limit of detection (copies/reaction), determining as the lowest concentration of the ten standard replicates that showed a standard deviation of $C_q$ of less than 0.5

[d] Assay limit of quantification (copies/reaction), determining as the lowest concentration of the target gene that can be correctly quantified

[e] From (Kongprajug et al., 2020)

**Table S4** Reproducibility of the qPCR methods using freshwater and seawater field duplicates

| Sampling site | HPyVs (log$_{10}$ copies/100 mL) | crAssphage (log$_{10}$ copies/100 mL) |
|---|---|---|
| MP1.1/1 | <ALOD[a] | 3.67 |
| MP1.2/1 | <ALOD | 3.59 |
| Mean | NA[b] | 3.63 |
| SD | NA | 0.05 |
| %CV[c] | NA | 1.49 |
| MP1.1/2 | 3.56 | 4.37 |
| MP1.2/2 | 3.03 | 4.26 |
| Mean | 3.29 | 4.31 |
| SD | 0.38 | 0.08 |
| %CV | 11.45 | 1.76 |
| MP1.1/3 | 2.85 | 3.41 |
| MP1.2/3 | 2.72 | 3.56 |
| Mean | 2.78 | 3.49 |
| SD | 0.09 | 0.11 |
| %CV | 3.23 | 3.13 |
| B.1-1 | 2.67 | <ALOD |
| B.2-1 | 2.02 | <ALOD |
| Mean | 2.35 | NA |
| SD | 0.46 | NA |
| %CV | 19.45 | NA |
| B1-5 | 3.47 | <ALOD |
| B2-5 | 2.95 | <ALOD |
| Mean | 3.21 | NA |
| SD | 0.37 | NA |
| %CV | 11.58 | NA |

| Sampling site | HPyVs ($log_{10}$ copies/100 mL) | crAssphage ($log_{10}$ copies/100 mL) |
|---|---|---|
| G1-2 | 2.76 | 3.20 |
| G2-2 | 2.61 | 3.01 |
| Mean | 2.68 | 3.11 |
| SD | 0.11 | 0.14 |
| %CV | 3.96 | 4.41 |
| G.1-4 | 3.00 | 2.55 |
| G.2-4 | 2.72 | 2.60 |
| Mean | 2.86 | 2.58 |
| SD | 0.20 | 0.03 |
| %CV | 7.09 | 1.28 |

[a]Assay limit of detection of 40 copies/reaction for the HPyV and 20 copies/reaction for the crAssphage assays
[b]Not available
[c]Coefficient of variation percentage

**Table S5** Qualitative MST marker distribution

| Sample | Assay (no. of positive samples/no. of total samples) | | | | |
|---|---|---|---|---|---|
| | HPyV | crAssphage[a] | Lachno3 | BacV6-21 | Positive in both Lachno3 and BacV6-21 |
| Human sewage | 19/19 | 19/19 | 19/19 | 18/19 | 18/19 |
| Swine | 0/38 | 0/38 | 27/37 | 20/38 | 14/37 |
| Cattle | 0/34 | 0/34 | 12/33 | 3/25 | 1/25 |
| Chicken | 0/23 | 0/23 | 1/23 | 4/23 | 1/23 |
| Duck | 0/5 | 0/5 | 0/5 | 2/5 | 0/5 |
| Goat | 0/10 | 0/10 | 6/10 | 0/10 | 0/10 |
| Sheep | 0/4 | 0/4 | 2/4 | 0/4 | 0/4 |
| Buffalo | 0/6 | 0/6 | 0/6 | 2/6 | 0/6 |
| TP[b] | 19 | 19 | 19 | 18 | 18 |
| FP[c] | 0 | 0 | 48 | 31 | 16 |
| TN[d] | 120 | 120 | 70 | 80 | 94 |
| FN[e] | 0 | 0 | 0 | 1 | 1 |

[a] Data for crAssphage were selected from (Kongprajug et al., 2019) for comparison using similar sample sets to this study

[b] TP (true positive) is the number of target samples that showed positive results

[c] FP (false positive) is the number of non-target samples that showed positive results

[d] TN (true negative) is the number of non-target samples that showed negative results

[e] FN (false negative) is the number of target samples that represented negative results

**Table S6** Abundance of genetic markers in human sewage and nonhuman fecal samples

| Assay | Sample | N[a] | No. of ND[b] | Min[c] | 25th percentile[c] | Median[c] | 75th percentile[c] | Max[c] |
|---|---|---|---|---|---|---|---|---|
| **HPyV** | **Sewage** | **19** | **0** | **3.66** | **4.93** | **5.31** | **5.49** | **6.53** |
| | Swine | 38 | 38 | <1.60[d] | <1.60 | <1.60 | <1.60 | <1.60 |
| | Cattle | 34 | 34 | <1.60 | <1.60 | <1.60 | <1.60 | <1.60 |
| | Chicken | 23 | 23 | <1.60 | <1.60 | <1.60 | <1.60 | <1.60 |
| | Duck | 5 | 5 | <1.60 | <1.60 | <1.60 | <1.60 | <1.60 |
| | Goat | 10 | 10 | <1.60 | <1.60 | <1.60 | <1.60 | <1.60 |
| | Sheep | 4 | 4 | <1.60 | <1.60 | <1.60 | <1.60 | <1.60 |
| | Buffalo | 6 | 6 | <1.60 | <1.60 | <1.60 | <1.60 | <1.60 |
| **crAssphage[e]** | **Sewage** | **19** | **0** | **5.28** | **6.07** | **6.36** | **6.77** | **7.38** |
| | Swine | 38 | 38 | <1.30 | <1.30 | <1.30 | <1.30 | <1.30 |
| | Cattle | 34 | 34 | <1.30 | <1.30 | <1.30 | <1.30 | <1.30 |
| | Chicken | 23 | 23 | <1.30 | <1.30 | <1.30 | <1.30 | <1.30 |
| | Duck | 5 | 5 | <1.30 | <1.30 | <1.30 | <1.30 | <1.30 |
| | Goat | 10 | 10 | <1.30 | <1.30 | <1.30 | <1.30 | <1.30 |
| | Sheep | 4 | 4 | <1.30 | <1.30 | <1.30 | <1.30 | <1.30 |
| | Buffalo | 6 | 6 | <1.30 | <1.30 | <1.30 | <1.30 | <1.30 |
| **Lachno3** | **Sewage** | **19** | **0** | **5.42** | **6.58** | **7.30** | **7.60** | **8.02** |
| | Swine | 37 | 10 | <1.70 | <1.70 | 7.67 | 8.53 | 9.12 |
| | Cattle | 33 | 21 | <1.70 | <1.70 | <1.70 | 5.16 | 7.45 |
| | Chicken | 23 | 22 | <1.70 | <1.70 | <1.70 | <1.70 | 8.21 |
| | Duck | 5 | 5 | <1.70 | <1.70 | <1.70 | <1.70 | <1.70 |
| | Goat | 10 | 4 | <1.70 | <1.70 | 6.49 | 6.68 | 6.87 |
| | Sheep | 4 | 2 | <1.70 | <1.70 | 6.54 | 6.99 | 6.99 |
| | Buffalo | 6 | 6 | <1.70 | <1.70 | <1.70 | <1.70 | <1.70 |

| Assay | Sample | N[a] | No. of ND[b] | Min[c] | 25th percentile[c] | Median[c] | 75th percentile[c] | Max[c] |
|---|---|---|---|---|---|---|---|---|
| **BacV6-21** | **Sewage** | **19** | **1** | **<1.70** | **6.66** | **7.28** | **7.70** | **8.05** |
| | Swine | 38 | 18 | <1.70 | <1.70 | 4.96 | 6.01 | 7.66 |
| | Cattle | 25 | 22 | <1.70 | <1.70 | <1.70 | <1.70 | 6.61 |
| | Chicken | 23 | 19 | <1.70 | <1.70 | <1.70 | <1.70 | 8.07 |
| | Duck | 5 | 3 | <1.70 | <1.70 | <1.70 | 5.26 | 7.04 |
| | Goat | 10 | 10 | <1.70 | <1.70 | <1.70 | <1.70 | <1.70 |
| | Sheep | 4 | 4 | <1.70 | <1.70 | <1.70 | <1.70 | <1.70 |
| | Buffalo | 6 | 4 | <1.70 | <1.70 | <1.70 | 6.17 | 7.13 |

[a] Total number of samples

[b] Total number of non-detected samples

[c] Genetic marker abundance in $log_{10}$ copies/g feces for nonhuman fecal samples, and in $log_{10}$ copies/100 mL sewage for human sewage samples.

[d] Not detectable (<ALOQ in $log_{10}$ copies/reaction)

[e] Data for crAssphage were selected from (Kongprajug et al., 2019) for comparison

**Table S7** Abundance of HPyV and crAssphage markers in municipal wastewater and environmental samples

| Assay | Sample | N[a] | No. of ND[b] | Min[c] | 25th percentile[c] | Median[c] | 75th percentile[c] | Max[c] |
|---|---|---|---|---|---|---|---|---|
| HPyV | WWTP1-inf | 6 | 0 | 4.38 | 4.54 | 5.39 | 5.78 | 6.03 |
| | WWTP1-eff | 6 | 0 | 3.91 | 4.31 | 4.71 | 4.85 | 4.86 |
| | WWTP2-inf | 6 | 0 | 5.69 | 5.73 | 5.78 | 5.97 | 6.02 |
| | WWTP2-eff | 6 | 1 | <1.70[d] | 3.17 | 3.45 | 3.67 | 5.09 |
| | Beach A | 17 | 1 | <1.70 | 2.31 | 2.74 | 3.09 | 4.12 |
| | Beach B | 24 | 2 | <1.70 | 2.33 | 2.49 | 2.74 | 4.33 |
| | Freshwater C | 27 | 2 | <1.70 | 3.04 | 3.45 | 4.41 | 5.10 |
| crAssphage | WWTP1-inf | 6 | 0 | 5.04 | 5.10 | 6.30 | 6.71 | 7.16 |
| | WWTP1-eff | 6 | 0 | 4.64 | 4.83 | 5.27 | 5.71 | 5.81 |
| | WWTP2-inf | 6 | 0 | 6.19 | 6.23 | 6.73 | 7.13 | 7.16 |
| | WWTP2-eff | 6 | 0 | 3.96 | 4.51 | 5.07 | 5.64 | 5.96 |
| | Beach A | 17 | 10 | <1.70 | <1.70 | <1.70 | 2.15 | 2.68 |
| | Beach B | 24 | 8 | <1.70 | <1.70 | 2.30 | 2.58 | 3.51 |
| | Freshwater C | 27 | 3 | <1.70 | 3.41 | 4.19 | 4.51 | 5.21 |

[a] Total number of samples

[b] Total number of non-detected samples

[c] Genetic marker abundance in $\log_{10}$ copies/g feces for nonhuman fecal samples, and in $\log_{10}$ copies/100 mL sewage for human sewage samples.

[d] Not detectable (<ALOQ in $\log_{10}$ copies/reaction)

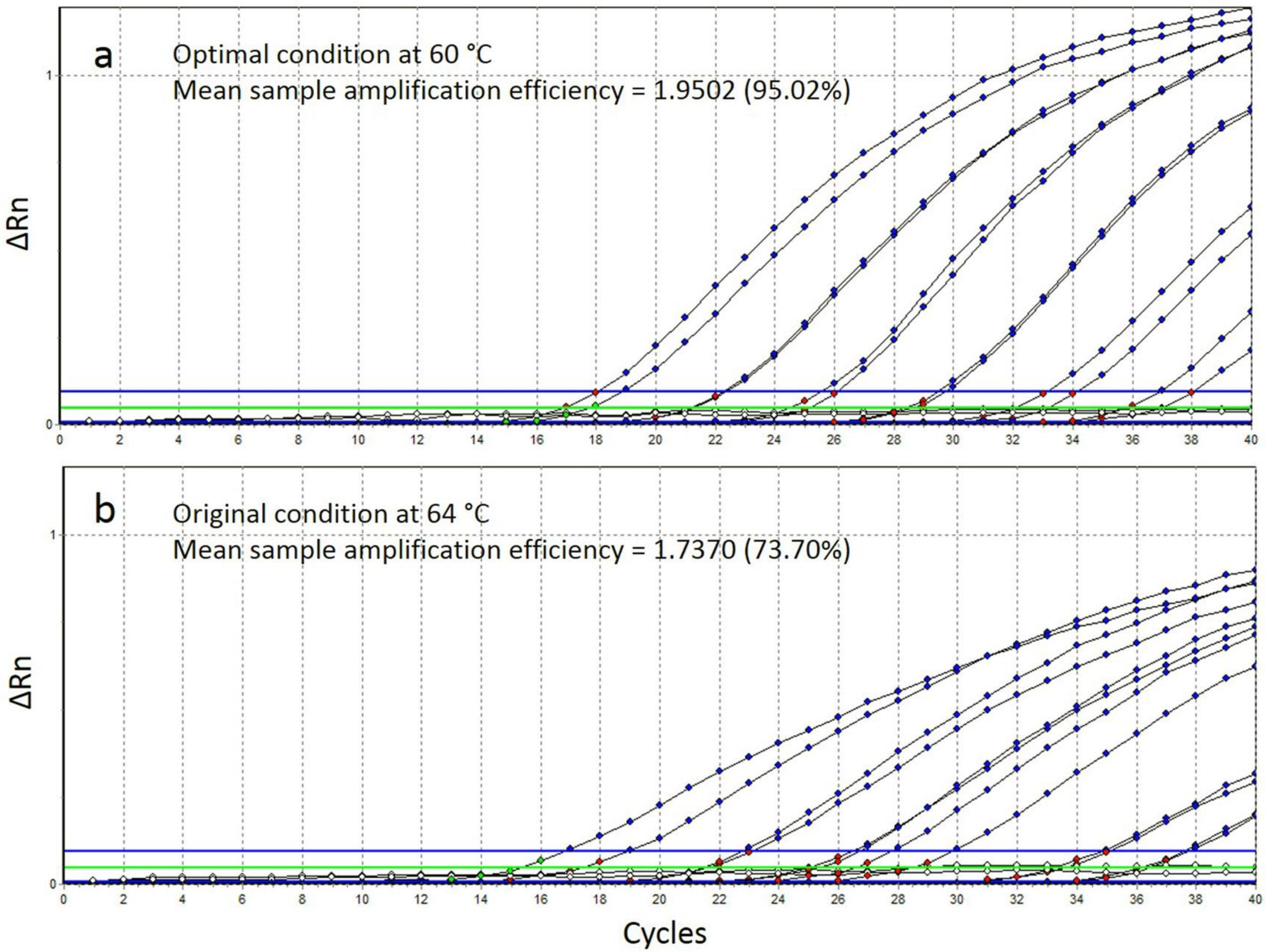

**Figure S1** Screenshots of the Lachno3 qPCR analysis with annealing temperatures of 60°C (a) and 64°C (b)
